# EMBRYONIC DEVELOPMENT OF THE FIRE-EYE-TETRA *Moenkhausia oligolepis* (CHARACIFORMES: CHARACIDAE)

**DOI:** 10.1101/2020.08.18.255554

**Authors:** Raquel Santos dos Santos, Jeane Rodrigues Rodrigues, Jhennifer Gomes Cordeiro, Hadda Tercya, Marissol Leite, Bruna Patrícia Dutra Costa, Raphael da Silva Costa, Caio Maximino, Diógenes Henrique de Siqueira-Silva

## Abstract

This study describes the embryonic development of *Moenkhausia oligolepis* in captive conditions. After fertilization, the embryos were collected every 10 min up to 2 h, every 20 min up to 4 h, and every 30 min until hatching. The fertilized eggs of *M. oligolepis* measured approximately 0.85 ± 0.5 mm and have an adhesive surface. The embryonic development lasted 14 hours at 25°C, with the Zygote, Cleavage, Blastula, Gastrula, Neurula and Segmentation phases. The hatching occurred in embryos around the 30-somites stage. Our results bring only the second description of embryonic development to a species of Moenkhausia genus, the first for the refereed species. Such data are of paramount importance considering the current conflicting state of this genus phylogenetic classification and may help taxonomic studies. Understand the biology of a species that is easily handling in captive conditions and has an ornamental appeal may assist studies in its reproduction in order to both, supply the aquarium market and help the species conservation in nature. Moreover, our data enable the *M. oligolepis* to be used as a model species in biotechnological applications, such germ cell transplantation approach.

## 1 INTRODUCTION

The study of embryological development is an important tool that allows the knowledge of a species life history (De Alexandre et al., 2009). This phase of development comprises fish formation events, from fertilization of the oocyte by spermatozoa to larval hatching (Solnica-Krezel, 2005). At this phase, the animal is more vulnerable to any environmental disturbance, which can change its morphology, cause deformities, or even the death. Therefore, in order to investigate the effects of changes in climatic variables on the embryonic development of teleosts, many studies describe this phase and associate its development with abiotic factors, such as temperature (Hansen and Peterson, 2001; Rodrigues-Galdino et al., 2010; Arashiro et al., 2018), water acidification (Villanueva et al., 2011), water dissolved O_2_ (Keckeis et al., 1996), among others.

Studies on embryonic developbment are also important to subsidize research on phylogeny and taxonomy of species, allowing the knowledge of evolutionary history and relations (Godinho and Lamas, 2009; Weber et al., 2012; Dos Santos et al., 2016). In addition, Godinho and Lamas (2009) showed that the characteristics of eggs, when fertilized, help in the knowledge of reproductive strategies of teleosts.

In Brazil, studies on embryology focus, mainly, on species in which a commercial value is already established, such as the Siluriformes *Pseudoplatystoma coruscans* (Cardoso et al., 1995; Marques et al., 2008), and the Characiformes *Colossoma macropomum* (Leite et al., 2013), *Brycon insignis* (Isaú et al., 2011), and *Brycon cephalus* (Romagosa et al., 2001; De Alexandre et al., 2009), among many other large sized animals. However, considering the abundance of described species, especially of freshwater fish (3,148 species described until 2018 - ICMBIO, 2018), those works evidently do not contemplate the diversity of species.

The genus *Moenkhausia* (Eigenmann, 1903), for example, covers about 90 species of freshwater fish distributed in South America: Venezuela, Guyana, Amazonia (Froese and Pauly, 2018), and all Brazilian watersheds (Lima and Toledo-Piza, 2001). This genus belongs to the Characiformes order and it is currently allocated as *Incertæ sedis* in Characidae family, due to the lack of detailed research about its phylogeny. Although some taxonomic studies have already been carried out (Hojo et al., 2004; Benine et al., 2007 and 2009; Carvalho et al., 2014), the current situation about its classification is still unclear, since most studies are limited to the description of species of the genus.

This is the case of the species *Moenkhausia oligolepis* (Gunther, 1864), which is currently undergoing discussions about its classification due to the wide distribution of *Moenkhausia* species, coexisting and exhibiting similarities of colors and patterns. For this reason, Costa (1994) and Benine (2009) propose *M. oligolepis* as a complex of species. However, according to Domingos et al., (2014), the coexistence and similarity between species usually results in an incorrect definition of their conservation status. Called in some areas as black tail tetra (Matos et al., 2003), this species achieves around 10 cm of total length, when mature (Froese and Pauly, 2018). Present a reticulated body color, reddish pigmentation on the dorsal margin of the eye, giving it popular name (fire-eye-tetra), and a dark spot on the stalks of the caudal fin.

Thus, in order to contribute to the knowledge about the biology and conservation of the species, besides helping to identify and classify it, this study aimed to describe the embryonic development of *M. oligolepis* under captive conditions. The study describes the timing of usual stages after fertilization, based on external morphology, in captive individuals of *M. oligolepis*. It was found that the embryonic development lasted 14 hours at 25°C, with staging occurring at similar times as that of closely related species (e.g., *Brycon gouldingi*: Faustino et al., 2011; *Astyanax bimaculatus:* Weber et al., 2012; *Astyanax altiparanae*: Dos Santos et al., 2016).

## 2 MATERIAL AND METHODOS

### Sampling of animals

The sexually mature individuals of *M. oligolepis* were collected in streams in the Tocantins Basin, located in the interior of the Amazon Forest, in the “Fundação Zoobotânica de Marabá” - PA (collection authorization ICMBio n° 62027-1). Nets (1.10 mm nylon, 4.75 x 1 mesh, 10 cm) were used to sample the fish, which were transported in 30-liter-plastic bags filled with water and equipped with portable aerator, to the laboratory. The species was identified in the Laboratory of Biology and Fish Genetics of the Institute of Biosciences of the Universidade Estadual Paulista (UNESP), Botucatu, state of São Paulo, Brazil (voucher: 25622).

Fish acclimatization lasted four months in glass tanks (23 × 21 cm, capacity of 13 liters of water) with aeration pumps and internal bacteriological filter. The animals were fed three times a day with commercial feed (4200 Kcal·kg^−1^ and 28% crude protein) and the tank water was partially exchanged daily.

### Preparation of matrices

Four males and three females were separated in a tank that had the same dimensions of the acclimatization ones, with constant circulation of water. Those animals were submitted to a monitored photoperiod cycle of 12 hours of light/dark, for 45 days. During this period, the water parameters (dissolved ammonia, nitrite, dissolved O_2_, pH and temperature) were analyzed every day. The same commercial feed was offered throughout the day in three plots of 0.100 g each, totaling 0.300 g of feed a day.

### Induction to spawning and fertilization

At the 45^th^ day, animals were injected with the pituitary crude extract of carp macerated and diluted in 0.9% saline solution. The solution was applied in the coelomic cavity at the base of the pectoral fin using an insulin syringe (1 ml) with a needle. Before this procedure, the animals were anesthetized with 1 ml of Eugenol solution (20 ml of Biodynamic Eugenol in 100 ml of Absolute Alcohol) diluted in 500 ml of water. This step was based on the protocol of Ninhaus-Silveira et al. (2006), in which females received two hormonal doses: the first doses of 0.5 mg / kg body weight and, after a 12-hour interval, the second doses of 5.0 mg / kg body weight. Males received a single dose of 1.0 mg / kg body weight at the same time as the second dose of females.

### Embryo collection and analyses

Samples were collected at the following time intervals after fertilization: every 10 min up to 2 hours per fertilization (hpf); every 20 min up to 4 hpf: and every 30 min until hatching. The sampled embryos were fixated in a solution of 2.5% glutaraldehyde sodium phosphate buffer 0.1 M, pH 7.3, and were observed using a trinocular stereoscope (TNE-10TN Opton). The images were captured using the TC Capture program and a digital camera (Samsung A3 2015(8mp) and processed by the CorelDRAW program (version 2018).

The embryonic development of *M. oligolepis* was classified in the standard phases (zygote, cleavage, blastula, gastrula, segmentation and hatching), based on previous studies Arashiro et al. (2018). The temperature and parameters of the water were monitored and documented during the development of the embryos.

## 3 RESULTS

### Egg sampling

The spawning occurred semi-naturally, two hours after the application of the last hormonal doses.

### Egg morphology

The fertilized eggs of *M. oligolepis* measure 0.85 ± 0.5 mm (mean±SD) in diameter. They are demersal, spherical and translucent after fertilization, and do not present oil drop. The chorion has an adhesive surface, and the perivitelline spaces measure 0.1± 0.02 mm (mean±SD) (Fig. 1).

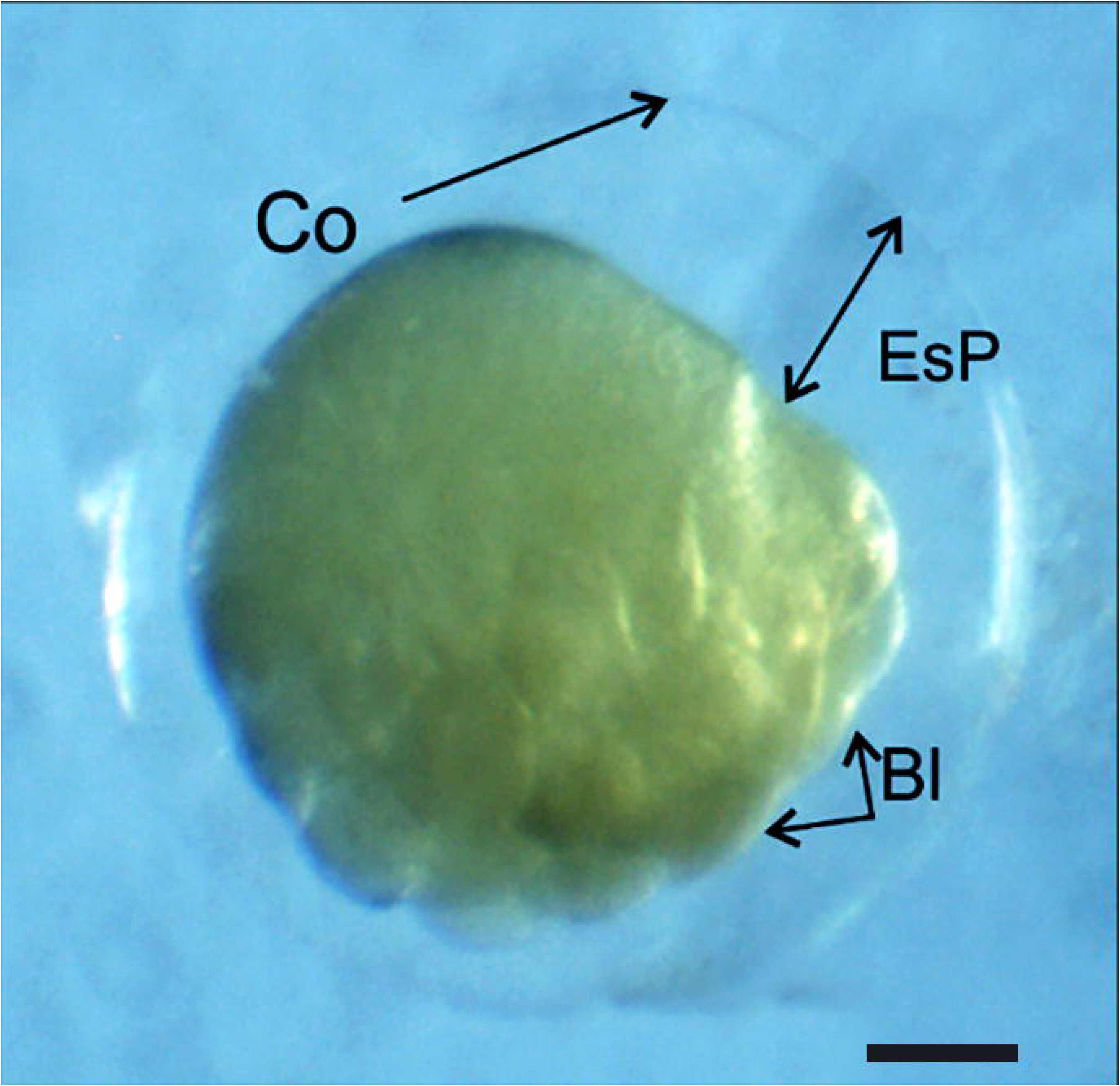

### Embryogenesis

Phases, stages and time of the development of *M. oligolepis* embryogenesis is listed on Table 1.

**Table 1.** Phases and time of embryonic development in *M. oligolepis* at the temperature of 25°C.

| **Phase** | **Stage** | **Time (h)** | **Fig.** |
| --- | --- | --- | --- |
| Zygote |  | 0.12 | 1A |
| Cleavage | 2 cell | 0.20 | 1B |
|  | 4 cell | 0.25 | 1C |
|  | 8 cell | 0.30 | 1D |
|  | 16 cell | 0.35 | 1E |
|  | 32 cell | 0.45 | 1F |
|  | 64 cell | 0.55 | 1G |
| Blastula | 128 cell | 1:10 | 1H |
|  | 256 cell | 1:20 | 1I |
|  | 512 cell | 1:30 | 1J |
|  | Dome | 1:40 | 1K |
| Gastrula | 50% epiboly | 2:00 | 1L |
|  | 75% epiboly | 2:40 | 1M |
|  | 90% epiboly | 3:30 | 1N |
|  | 95% / epiboly | 4:00 | 1O |
|  | Initial neurula | 4: 20 | 1P |
| Neurula | 100% epiboly | 5:30 | 2A |
| Segmentation | 5 somites | 5:50 | 2B |
|  | 8 somites | 6:30 | 2C |
|  | 17 somites | 9:30 | 2D |
|  | 27 somites | 11:30 | 2E |
| Hatching | 30 somites | 14:00 | 2F |

#### Zygote phase

It was observed an increase of the perivitelline space, and the formation of the blastodisc defining the animal and vegetal poles and evidencing a great quantity of yolk (Fig. 2A).

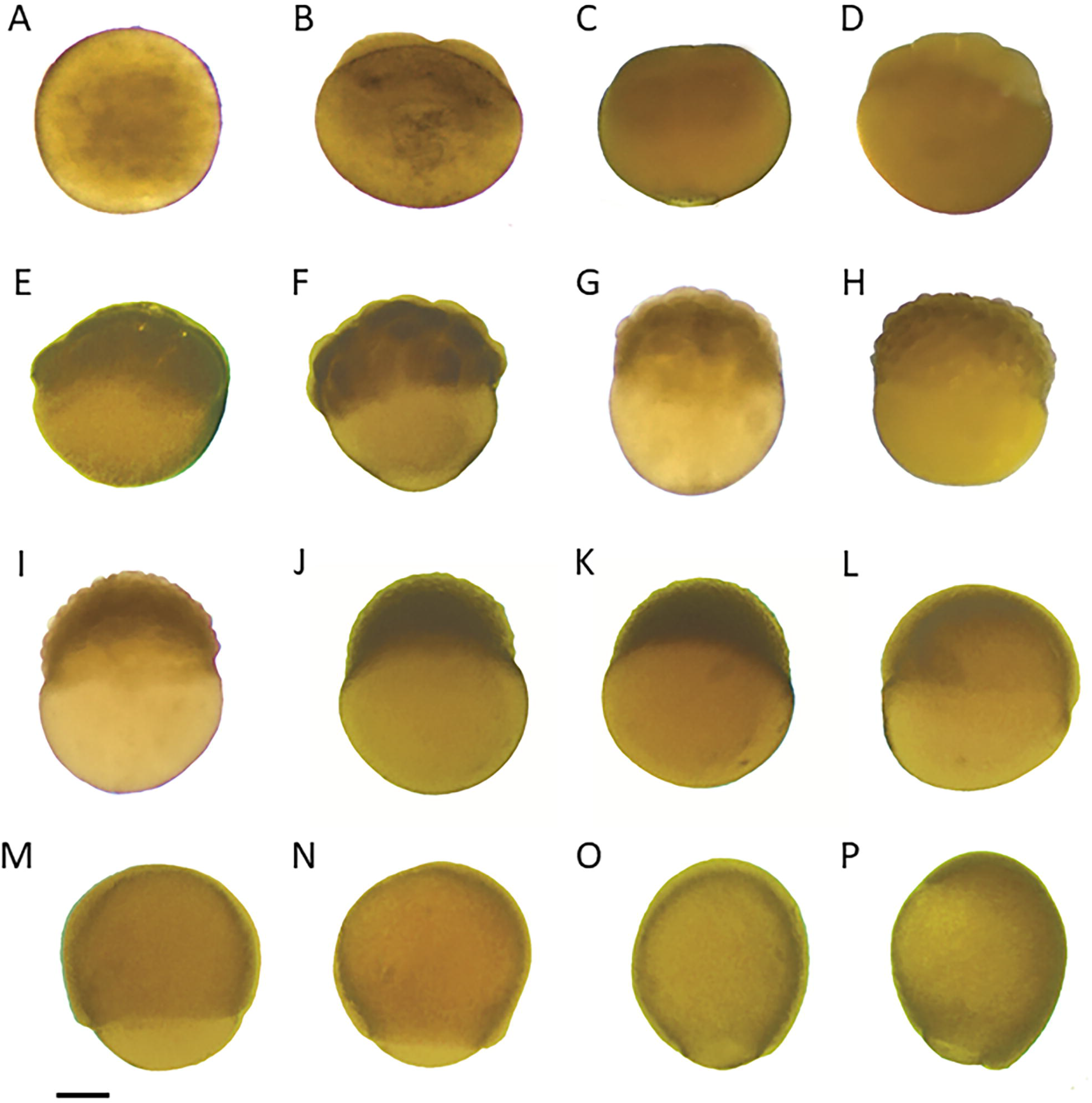

#### Cleavage phase

Cleavage followed the pattern of discoidal meroblastic division, being observed the presence of 2, 4, 8, 16, 32 and 64 consecutive blastomers (Fig. 2B-G). This phase took approximately 30 minutes.

#### Blastula phase

This phase was initiated at the sixth cleavage, doubling the number of cells in the sequences of 128, 256, and 512 blastomeres achieved at 1 h 30 min after fertilization (AF). The dome phase was reached at 1 h 40 min AF characterized by the organization of thousands of blastomeres in several layers at the top of the yolk, presenting a similar appearance to a mulberry (Fig. 2H-K).

#### Gastrula phase

This phase began around 2 h AF. The cells of blastoderm started the epiboly movement, moving toward the yolk and gradually evolving. At 2 h 40 min, a germinative ring was observed (Fig. 2L), and at 4 h AF 90% of the yolk was surrounded by the blastula and the blastopore was observed (fig. 2L-R).

#### Neurula

This stage occurred at 5 h and 30 min AF characterized mainly by epiboly of 100% of the embryo, whose blastoderm completely involves the yolk through epiboly (Fig. 3A).

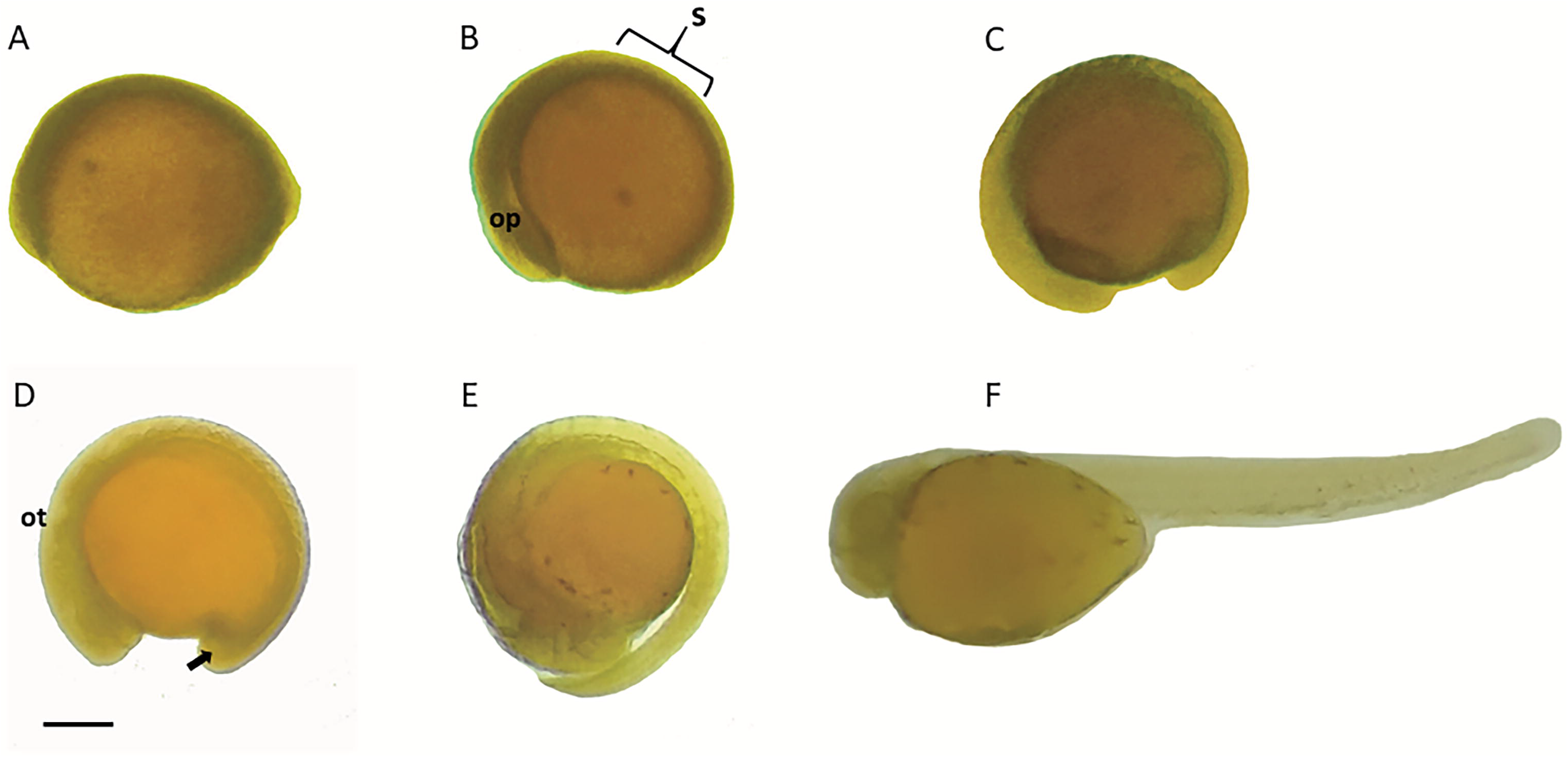

#### Segmentation

The segmentation phase is the last phase of embryonic development, and represents the differentiation of the cephalic and caudal poles, and the appearance of somites, vesicles and some external and internal organs of the embryo, extending until the moment of hatching. Segmentation lasted about 8 h 50 min. The embryo presented the first somite around 5 h 50 min AF, eight somites at 6 h30 min, and at 7 h30 min it was possible to visualize the optical vesicle. At 7 h40 min AF there were 17 somites, 8 h 10 min AF the appearance of the Kupffer vesicle was observed, followed by the appearance of the otic vesicle at 9 h AF. At 11 h 30 min AF, there were 27 somites, and after that, just before hatching, about 30 somites (Fig. 3B-F).

#### Hatching phase

The embryo presented free tail at 12 h 30 min AF, followed by larvae hatching at 14 h AF, with about 30 somites (Fig. 3F).

## 4 DISCUSSION

In this study we described the embryonic development of *M. oligolepis*, a Characidae of disputed taxonomic position from the Amazon, up to hatching. We found that the embryonic development lasted 14 hours at 25°C, with staging occurring at similar times as that of closely related species.

The ontogenetic development in fish is sensitive to changes in temperature, since its metabolic activities can be accelerated or retarded, altering the rhythm of the embryonic development (Santos et al., 2006; Faustino et al., 2010). This period is variable among species, and may be short as observed in *M. oligolepis*, or even shorter, as in *M. sanctaefilomenae*, whose embryonic development lasted 13 h (Walter, 2011). On the other hand, *Prochilodus lineatus* at higher temperatures (28°C) presented embryo development time similar to that of the present study (Ninhaus-Silveira et al., 2006), which makes clear that each species has its own relation with abiotic factors, a strategy that reflects the life history of each species.

The diameter of eggs is also directly related to the reproductive strategy, since small eggs are usually found in migratory species of total spawning and the largest in non-migratory species (Godinho et al., 2010). The diameter of *M. oligolepis* eggs is similar to those observed by Sato et al., (2006) and Webber et al., (2012) in other small Characiformes, *Astyanax bimaculatus* and *Tetragonopterus chalceus,* respectively, both reofilic species. *Astyanax bimaculatus* also reproduces in lentic waters (Webber et al., 2012).

It was also observed that the eggs of *M. oligolepsis* show characteristics of adhesiveness. According to Kolm and Ahnesjö (2005), adhesive eggs are a characteristic of the species with partial spawning and parental care. Godinho et al., (2010) also observed higher adhesiveness in eggs of lentic species with multiple spawning, whereas lotic species presented free eggs and total spawning. Judging from the characteristics of the environment in which the matrices of this study were collected, we can suggest that *M. oligolepis* is a species that spawns in lentic waters; however, differently from other species with adhesive eggs, there is no evidence of parental care in this *M. oligolepis*.

The adhesion of the eggs to the substrate contributes to the viability and protection of the offspring in the natural environment, but in captivity it may cause great mortality of the embryos, since the eggs and embryos agglomerate impairing the gas exchange between the developing embryo and the external environment. Moreover, egg adhesion can contribute to the proliferation of fungi and bacteria, causing death or malformation in the embryos. Many techniques have been developed to mitigate such damages (Siddique et al., 2014), such as incubators equipped with a closed water recirculation system, which promote the circulation of water and embryos, preventing their deposit and agglomeration in the tank bottom (Luz et at., 2001). In the studied species, however, it was observed that, although the eggs presented strong adhesiveness forming clusters of embryos, fixed to each other, or to the walls of the aquarium, the aerator was sufficient to keep them suspended in the water, dispensing the use of more elaborate techniques.

Another important structure in the embryological staging of fish is the chorion, since with the hydration of the egg it expands to form the perivitelline space (Siddique et al., 2014), which will aid in the development of the embryo, protecting it from external injuries often caused by the water flow. Due to this, eggs with large perivitelline space are characteristic of species that reproduce in agitated waters, whereas, smaller spaces are present in eggs of species that spawn in calm waters, an aspect that reflects different adaptations of the species to the environment they live (Yamagami et al., 1992; De Alexandre et al., 2009; Ribeiro et al., 2012; Yang et al., 2014). Similar to other Characiformes, such as *Acestrorhynchus spp., Hoplias lacerdae, Prochilodus spp., Leporinus sp.* observed by Rizzo et al., (2002), *M. oligolepis* presents pelagic eggs.

Considering that this is only the second embryological study of the genus *Moenkhausia*, this work brings important data about the embryology of *M. oligolepis*. We note that although much information has been revealed and supported, some of them need more detailed and elaborate assessments and we encourage the use of such data to clarify the confusing picture in species and genus classification. As suggested by Webber et al., (2012), studies like this are important to support future studies on reproduction, phylogeny and taxonomy.

## Acknowledgments

We thank the Laboratory of Neurosciences and Behavior “Frederico Guilherme Graeff” (LaNec) for the structure used. The authors also thank the prof. Claudio de Oliveira from the Laboratory of Biology and Fish Genetics of the Institute of Biosciences of the State University of São Paulo (UNESP), for the identification of fish. To the Laboratory of Science and Technology of Madeira (UEPA) in the person of prof. Luiz Eduardo de Lima Melo, for facilities support. To all the students of the Research Group of Studies in Reproduction of Amazonian Fish (GERPA) and to the Fundação Zoobotânica of Marabá - PA for allowing the collection of fish in its facilities.

## Ethical standards

The authors assert that all procedures contributing to this work comply with the ethical standards of the relevant national and institutional guides on the care and use of laboratory animals.

